# Open Access and Altmetrics in the pandemic age: Forescast analysis on COVID-19 literature

**DOI:** 10.1101/2020.04.23.057307

**Authors:** Daniel Torres-Salinas, Nicolas Robinson-Garcia, Pedro A. Castillo-Valdivieso

## Abstract

We present an analysis on the uptake of open access on COVID-19 related literature as well as the social media attention they gather when compared with non OA papers. We use a dataset of publications curated by Dimensions and analyze articles and preprints. Our sample includes 11,686 publications of which 67.5% are openly accessible. OA publications tend to receive the largest share of social media attention as measured by the Altmetric Attention Score. 37.6% of OA publications are bronze, which means toll journals are providing free access. MedRxiv contributes to 36.3% of documents in repositories but papers in BiorXiv exhibit on average higher AAS. We predict the growth of COVID-19 literature in the following 30 days estimating ARIMA models for the overall publications set, OA vs. non OA and by location of the document (repository vs. journal). We estimate that COVID-19 publications will double in the next 20 days, but non OA publications will grow at a higher rate than OA publications. We conclude by discussing the implications of such findings on the dissemination and communication of research findings to mitigate the coronavirus outbreak.

## 1 Introduction

On March 11, 2020, the World Health Organization (WHO) declared the COVID-19 a world pandemic (Organization et al., 2020). Since then, the spread of the disease has expanded, forcing governments to confine their population and enforce social distancing to reduce the spread of the virus. The gravity of the situation has led to an unprecedented scientific race to mitigate the effects of the pandemic (Torres-Salinas et al., 2020) which is has overflowed the scientific scholarly communication system (Larriviere et al., 2020). The normal pace of scholarly communication has proven to be too slow and inefficient, leading to a complete transformation in the way new findings are reported and consumed. Traditional bibliometric databases such as Web of Science or Scopus, which index mainly published journal literature, have become almost instantly obsolete while journals are accelerating to an unprecedented rate their publication track for any COVID-related study. This has led scientists’ attention to unexpected sources such as *ad hoc* compilations of scientific literature openly accessible and curated by the scientific community. Examples of such compilations are the CORD-19 dataset^1^, a global community effort, the COVID-19 Database maintained by the WHO^2^, or some publisher curated lists. These topic-specific databases are characterized by their daily update as well as including both, peer-reviewed and non-peer reviewed literature literature.

The COVID-19 pandemic has confronted scientists to an unprecedented challenge in which time and efficiency are critical. The exponential growth of scientific literature on the coronavirus outbreak and the means by which new findings are disseminated, disregarding the traditional status of journals for the sake of speed and efficiency (Larrivière et al., 2020), confront scientists to additional obstacles. They need to keep up with new scientific literature, be more critical than ever with non-peer-reviewed literature and respond to the the expectations of society. As a consequence, scientific discussions and conflicts are more public than ever (Gulbrandsen et al., 2020), revealing an additional threat, as socially responsible attitudes are crucial to stop the spread of the outbreak (Thelwall and Thelwall, 2020). Examples such as a recent paper suggesting the virus was man-made (Delgado López-Cózar et al., 2020) reveal that responsible communication to non-scientific audiences is essential to balance between open scientific debates and public outreach. In this new context, altmetrics gain more importance than ever, as they become the quickest vehicle to monitor social perception of science in an area where citations play a secondary role as they lack on speed to keep up the production and reception of new findings.

In this study we compare the growth on publications, citations and altmetric mentions to COVID-19 literature using the Dimensions dataset which includes publications, datasets, grants and clinical trials (Resources, 2020).

The general goal is to analyze the size of scientific literature is expected in relation to this crisis, as well as the size of the discussions as any type of analysis or tool built based on this increasing body of information will have to consider such growth rate. More specifically, in this study we aim at responding at the following research questions:

1. What are the differences in terms of access to COVID-19 related literature? We establish comparisons between OA and non-OA output as well as between journal articles and preprints to study the effectiveness of the communication strategies followed by scientists working on this subject.
2. What is the expected growth of both, scientific literature, citations and social media attention? By modelling our data we establish predictions to up to 30 days which will can help on the design of infrastructure and tools which will make use of this data.

## 2 Data and methods

### 2.1 Data collection

We use the Dimensions dataset on COVID-19 literature version 14, which was updated for the last time in April 14, 2020 (Resources, 2020). This dataset contains information on four document types: publications, datasets, clinical trials and grants. In this study we work with the publications dataset, which includes a total of 11,686 records. This dataset is much more restrictive than CORD-19, which employs a much wider criteria of inclusion (Colavizza et al., 2020). This set is retrieved from the Dimensions database after using the following search query^3^:

Year: 2020; Data Search: “2019-nCoV” or “COVID-19” or “SARS-CoV-2” or ((“coronavirus” or “corona virus”) and (Wuhan or China))

For each record it includes publication metadata as well as information on number of citations, Altmetric Attention Score, journal or repository and open access (OA) status. Table 1 offers a brief overview of the contents of the publication dataset with regard to publication type, document type and type of access. Dimensions provided OA information retrieved from Unpaywall, but assigns documents to one OA type exclusively, overriding cases in which there might be evidence of more than one OA type for a single document (Robinson-Garcia et al., 2020).

**Table 1.** Overview of the Dimensions dataset for COVID-19 related publications

| Type of access | Journal | Repository | % preprints |
| --- | --- | --- | --- |
| Closed | 3514 | 288 | 0.00 |
| Bronze | 3072 | 318 | 0.00 |
| Green, Accepted | 4 | 0 | 0.00 |
| Green, Published | 15 | 627 | 0.98 |
| Green, Submitted | 21 | 1538 | 0.99 |
| Hybrid | 458 | 626 | 0.00 |
| Pure Gold | 1205 | 0 | 0.00 |
| **Total** | **8289** | **3397** | **0.23** |

In this study we restrict our analysis to two document types, that is, articles and preprints. The Dimensions dataset includes other document types such as monographs, book chapters and proceedings, but these only amount to a total of 278 records. We must also note that preprints and articles are document types unrelated to their OA status of the manuscript, as preprints (e.g., online first) can also be found in journals. After some normalization on the original dataset, we identified 67.5% of all COVID-19 related publications openly accessible, with 8.2% of closed publications deposited under embargo in repositories.

### 2.2 Methods

The focus of the paper is on the growth of publications as well as of social media attention. As a proxy for the latter we use the Altmetric Attention Score (AAS) provided in the original dataset. Altmetric scores can only be obtained for documents which include an identifier such as a DOI or a PMID. 11,189 records in the Dimensions dataset include an identifier, that is 95.7% of the records. The AAS has been strongly criticized by the scientometric community (Gumpenberger et al., 2016; Mukherjee et al., 2018) as it is a composite measure difficult to interpret. In the case of altmetrics this becomes even more problematic as Altmetric.com (the altmetric platform behind the score) includes a plethora of diverse sources with little relation with each other. While these limitations are acknowledged, we used this indicator as an exploratory attempt to identify those documents with higher social media attention. In further analyses we plan to obtain additional information from Altmetric.com on the specific scores obtained by each paper in each of the platforms this database covers.

To establish prediction on publications, citations and altmetrics growth (with particular interest on OA) we address the proble as a one one of time series prediction. To do so, we need adequate tools to analyze historical data and thus, making predictions Hassan (2014); de Oliveira and Oliveira (2018). There are several types of models that can be used for time-series forecasting Siami-Namini et al. (2018). In this study we make use of ARIMA (AutoRegressive Integrated Moving Average) Ho and Xie (1998), which is one of the most widely known approaches Hyndman and Athanasopoulos (2018). In this kind of models, the forecasts correspond to a linear combination of past values of the variable Hyndman and Khandakar (2008), explaining a given time series based on past values.

An ARIMA model is characterized by three parameters (*p, d, q*) where,

– *p* refers to the use of past values in the regression equation for the series
– *d* indicates the order of difference for attaining stationarity
– *q* determines the number of terms to include in the model

Here we obtain ARIMA models for the total number of publications, by location of the record (journal or repository) and OA status. All the analyses are conducted on an Ubuntu 18.04.1 machine with R version 3.6.3 and RStudio version 1.1.456. Figure 1 shows the publication time trends observed in the Dimensions dataset. As reported in a previous paper (Torres-Salinas, 2020) the literature on COVID-19 is growing at an exponential rate. If we consider the total number of publications, the value of *R*^2^ is equal to 0.93. In the case of journal publications the value of *R*^2^ is 0.92. In the case of repositories, we observe a much slower growth (*R*^2^ = 0.36). Predictive models were obtained for each of the variables observed and subsequently estimated. These models will be referred to from here on as ARIMA(1,2,2) for the “‘Total” series, ARIMA(0,2,1) for the “Journal” series, and ARIMA(2,2,4) for the “repositories” series. Our 30 days predictions are based on these models.

**Fig. 1.**
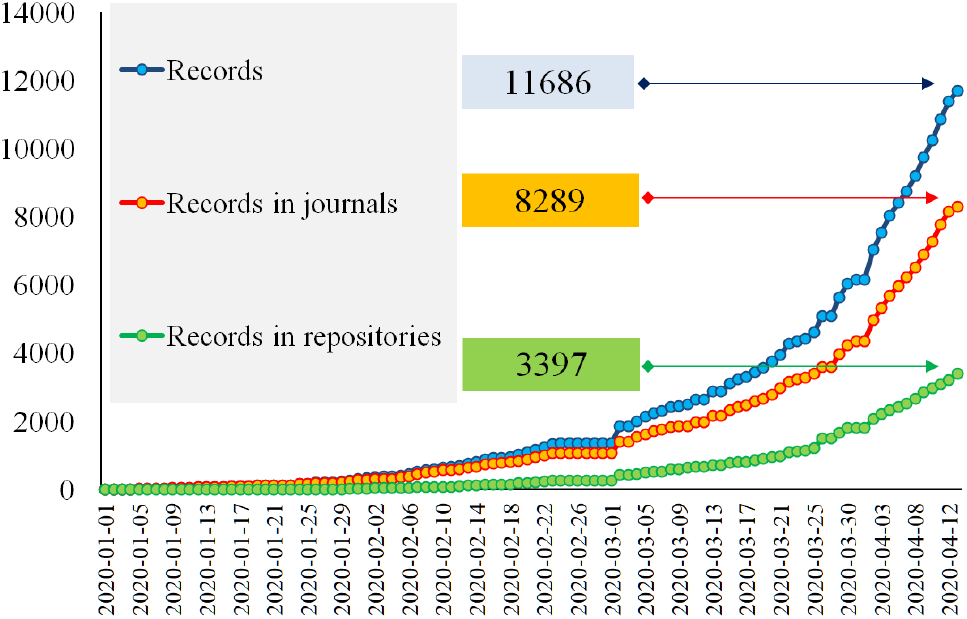
Time trend on the accumulated number of records overall, in journals and in repositories

## 3 Results

### 3.1 Descriptive analysis

Table 2 provides an overview of the dataset used. A total of 11,686 papers were retrieved, out of which 7,884 (68%) are available in OA. This proportion decreases during the month of April. Despite the fact that this analysis covers three and 1/2 months, a total of 27,129 citations have already been made. This means on average 2.32 citations per paper. This average is even higher for non OA publications, which receive an average number of citations of 3.28.

**Table 2.** Description of the dataset by type of access. **A** Time trend, **B** Altmetric Attention Score and citation indicators, and **C** distribution of records, Altmetric Attention Score and citations by location (journals or repositories). * These two repositories include also journal literature and hence overlap with the two location types.

| a. Number of papers in *Dimensions* |  |  |  |
| --- | --- | --- | --- |
|  | Totals | Open Access | Non Open Access |
| ▪ January | 313 | 261 | 52 |
| ▪ February | 1039 | 847 | 192 |
| ▪ March | 4815 | 3980 | 835 |
| ▪ April (until 04/13) | 5519 | 2796 | 2723 |
|  | 11686 | 7884 (68%) | 3802 (32%) |
| b. *Attention Altmetric Score* & citations |  |  |  |
|  | Accumulated | Average per paper | Max Value |
| *Altmetric Attention Score- AAS* | 1373008 | 117.49 | 27609 |
| ▪ Open Access | 1200466 | 152.25 | 27609 |
| ▪ Non Open Access | 172542 | 45.38 | 7680 |
| *Citations* | 27129 | 2.32 | 1238 |
| ▪ Open Access | 25914 | 0.31 | 1238 |
| ▪ Non Open Access | 1215 | 3.28 | 127 |
| c. Journals, repositories and main information sources |  |  |  |
|  | Nr Papers | AAS accumulated | Citations accumulated |
| Journals | 8288 | 1105043 | 24604 |
| Repositories | 3397 | 267964 | 2525 |
| ▪ *Information sources:* |  |  |  |
| ▪ Pubmed* | 5143 | 998682 | 24008 |
| ▪ PMC* | 2330 | 424181 | 13973 |
| ▪ BioRxiv | 346 | 77411 | 1029 |
| ▪ MedRxiv | 1232 | 154540 | 1260 |

These papers have raised an unprecedented amount of social media attention according to their AAS. On average, these documents receive an AAS of 117, which is even higher in the case of OA papers (152.25). In this sense, we observe differences depending on the location of the record. Journal articles receive higher citations than those stored in repositories, but there are differences by repository. PubMed and PMC receive a considerably higher number of social media attention than the rest of the repositories. Although BioRvix and MedRxiv provide a lower number of documents to the dataset, they still attract a high number of citations (in the case of the former) and social media attention (for the latter).

### 3.2 Open Access and social media attention

Figure 2 shows the distribution of AAS (A) and numberthe relation between the number of documents and AAS each receives (B) by OA status. Most of the papers on COVID-19 are OA and reach higher values of AAS than non-OA papers. For instance, two papers obtained AAS values of 27,609 (Nat Med 26, 450-452, 2020) and 21,738 (N Engl J Med, 382, 1564-1567, 2020) respectively. Likewise 15 OA papers obtained a score of at least 10,000 AAS, accumulating a total of 215,885. These fifteen papers alone add up to more AAS than the entire set of papers published as non OA (Table 2).

**Fig. 2.**
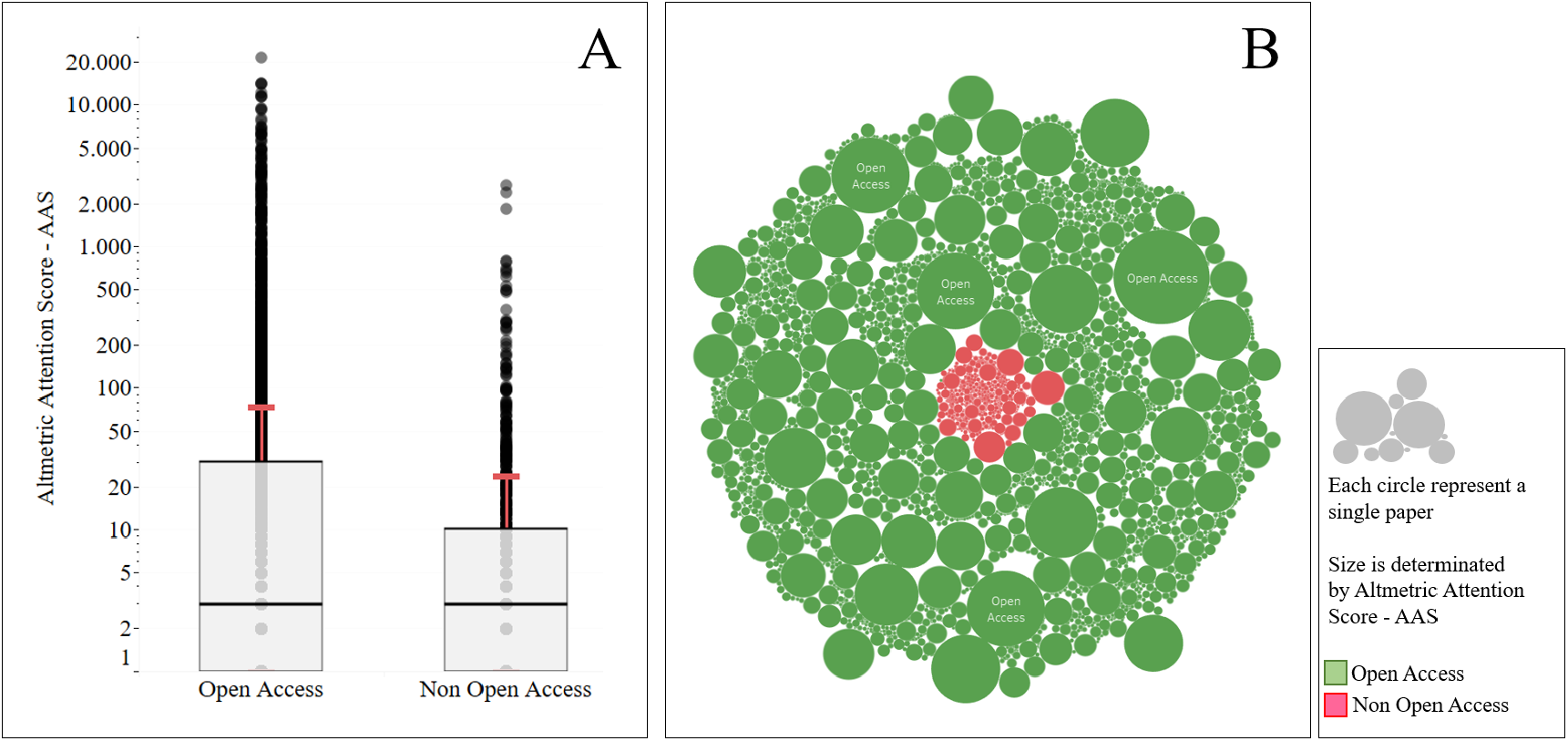
Altmetric Attention Score: Open Access and Non Open Access

Figure 3 shows the distribution of AAS (A) and the size of the output (B) by OA type. Bronze OA documents tend to receive a higher AAS and represent the largest share of COVID-19 related publications (3,072). Overall OA papers, either We observe that OA papers published in journals (regardless of the OA type: bronze, hybrid or pure), predominate. In relation to AAS, bronze papers have an average of 249 and papers with higher AAS are within this modality. Hybrid and gold OA papers receive less attention, 154 and 61 on average, respectively.

**Fig. 3.**
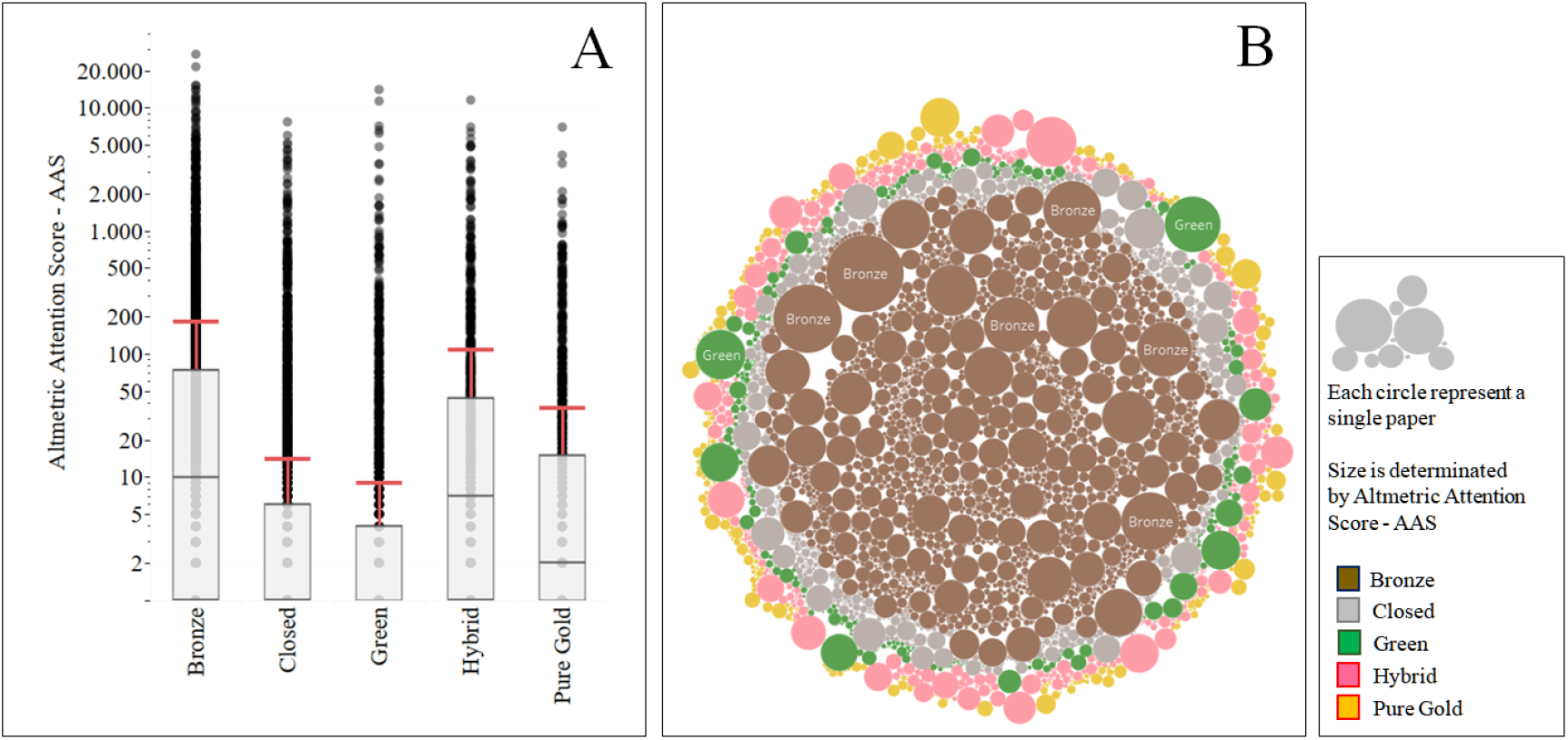
Overview of social media attention on open access. **A** Altmetric Attention Score distribution by OA type and **B** Number of records and AAS received by paper by OA type

In Figure 4 we shift our focus to records deposited in repositories. The repository with the largest number of publications is PMC, with a total of 2,330 papers and an average AAS of 182. Here we must note that PMC not only includes self-archived documents, but also indexes OA journals (Robinson-Garcia et al., 2020). The second largest repository is medRxiv with a total of 1,232 and an average AAS of 125 per document. Despite being the repository with the lowest number of records included (387), documents indexed in BioRxiv receive on average, the highest AAS (223). All documents in BioRxiv have receive at least an AAS of 1. The rest of the repositories analyzed (ChemRxiv, JMIR Preprints, Research Square and SSRN) have a peripheral role on production and visibility of COVID-19 related publications.

**Fig. 4.**
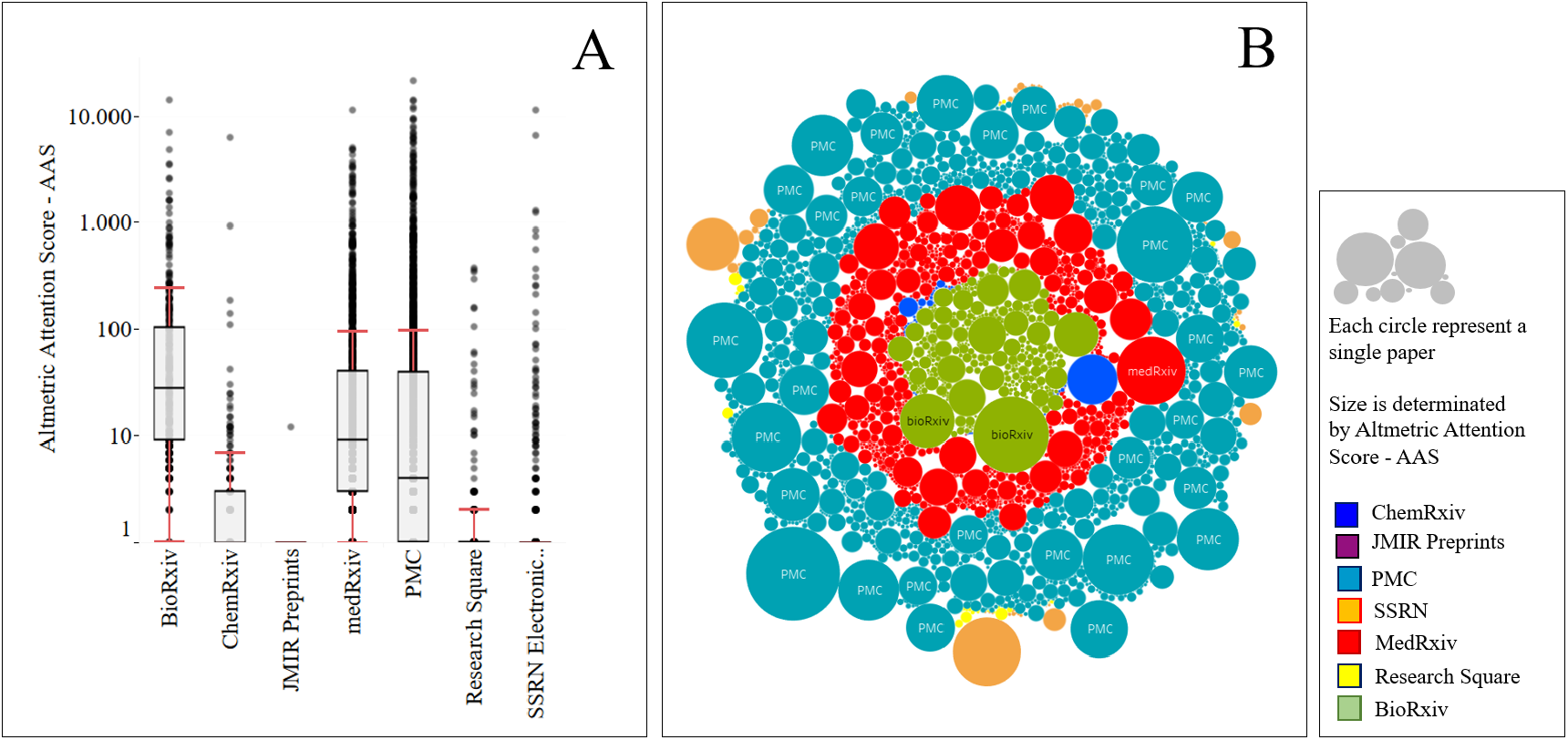
Overview of social media attention for documents deposited in repositories. **A** Altmetric Attention Score distribution by repository and **B** Number of records and AAS received by paper by repository

### 3.3 Predictive analysis: ARIMA models

Figure 5 shows the accumulated time trend on number of publications and by journal and repository, as well as the predicted trends according to the obtained ARIMA model.

**Fig. 5.**
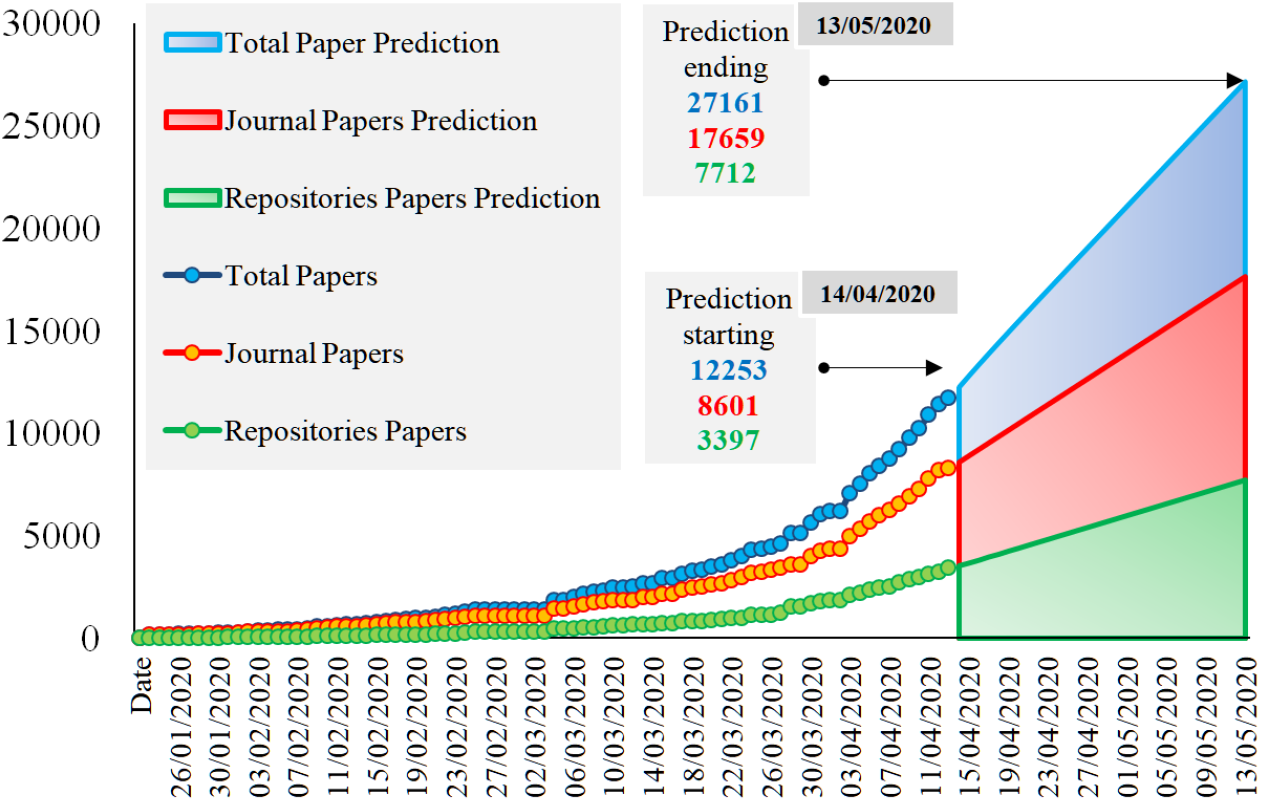
Growth evolution and predicted trend on COVID-19 related literature and by location (journals and repositories)

Paying attention to Figure 5, it can be seen that the estimate made by the predictive models at 30 days is of a growing trend in the number of publications. The ARIMA model forecast for total publications starts on 14/04/200 with 12254 publications and ends on 13/05/2020 with 27162 publications. In the case of journals it starts with 8601 publications and ends with 17660. The repositories will grow at a slower rate, the forecast starts with 3538 publications and ends with 7712. The data indicate that total publications will double in about 20 days, journal publications will double in about 24 days, and repository publications will double in about 24 days

Figure 6 shows the accumulated time trend as well as the predicted trend differentiating between OA and non OA publications.The ARIMA model forecast for OA papers starts on April 14, 2020 with 8,067 publications and ends on May 13, 2020 with 13,359 publications. According to these predictions, non OA publications will grow at faster rate than OA publications. The forecast starts with 4,075 publications and ends with 11,992. It can be said that OA publications will double every 30 days and non OA publications will double every 14 days. The differences between the number of OA and non OA publications appears to be narrowing as the prediction progresses. By the end of the forecast, the central role of open access will not be as clear as it was in early February and March.

**Fig. 6.**
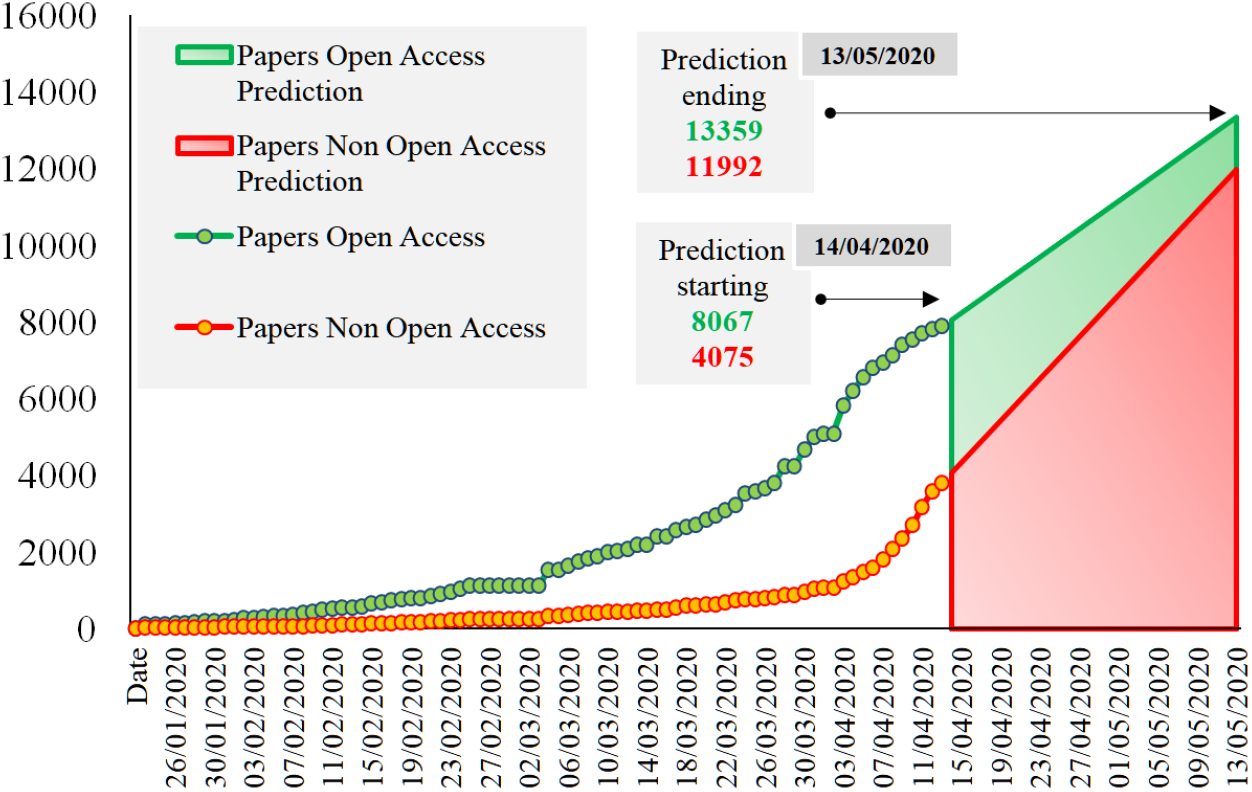
Growth evolution and predicted trend on COVID-19 related literature by OA and non OA.

## 4 Discussion and further research

This paper reports on the growth of scientific literature, citations and social media attention revolving around COVID-19 literature. For this, it uses the Dimensions dataset (Resources, 2020) which is openly accessible and has been updated daily until its last update on April 14, 2020. While the dataset itself is not free of limitations, and other COVID-19 datasets are being used alternatively, it is the one coming from the largest scientific database as compared with Web of Science and Scopus (Torres-Salinas et al., 2020). Furthermore, the search query used seems to be much more restrictive than other used elsewhere, which can introduce some noise when identifying the scientific corpus specifically dealing with this virus (Colavizza et al., 2020).

The findings reported here shows that many journals (e.g., New England Journal of Medicine, The Lancet, JAMA, Nature) are doing an important effort to prioritize the urgency of the current situation over their monetary benefits by providing COVID-19 related literature in OA. This is an unprecedented event which should not go unnoticed, and explains to a large extent the large shares of OA literature identified related with the coronavirus outbreak. The interest on scientific development on this front go beyond the scientific realm as the high social media attention revolving these documents shows. Scientific advancements are reported daily in the news media, discussed on Twitter and used for decision-making by politicians. Indeed, scientific efforts have not only focused on mitigating the pandemic, but have also responded to social concerns, such as those derived from the rise of fake news (Andersen et al., 2020).

The amount of literature produced since the coronavirus outbreak suggests an exponential growth on the number of publications produced, citations and social media mentions. If we want to be able to keep up with such growth and produce tools and analyses on such increasing corpus, some preparation is needed. Our estimates indicate that this number of records will duplicate every 14 days if the current rhythm of production continues. First reactions praised OA efforts from the scientific community and how these confronted the “normal” speed of science (Larrivière et al., 2020). However, our analysis shows a great dependency on journal literature and specifically on the role of major toll journals which have made openly accessible COVID-19 related literature as an exceptional measure. This reflects a great dependency on the traditional journal publishing system. Furthermore, our predictions estimate a higher growth for non OA literature in the near future. This trend, if confirmed, can become a great obstacle on the advancement of a cure for COVID-19 as well as on mitigating collateral damages from the pandemic.

Social media attention revolves mainly around OA publications, but again, here the role of toll journals opening their contents through bronze OA is crucial, followed by hybrid OA and gold OA, again reflecting that, despite the urgency, the traditional and mechanisms of scholarly publishing are still in place, along with all their deficiencies (Gadd, 2020).

That said, any conclusions on the predictions reported must be taken with caution as we live in a constantly changing situation, closely linked to the mitigation of the pandemic and political actions derived from it. Still, analyses such as the present can help us contextualize the phenomenon and provide alternative views from which scientometricians can contribute.

## 5 Summary of key findings

In this section we provide a brief summary of the main findings reported in this study.

1. 11,686 publications on COVID-19 were retrieved. 68% are OA. 27,129 citations have already been made. This means on average 2.32 citations per publication
2. On average publications receive an Altmetric Attention Score of 117, which is even higher in the case of Open Access papers (152.25)
3. Most of the publications on COVID-19 are OA and receive higher social media attention than non OA papers.
4. Most of the OA publications are bronze OA. These are receiving the highest social media.
5. OA papers published in scientific journals predominate. This fact emphasizes the central role of journals and peer review versus early access to preprints.
6. Pubmed is the repository with the largest number of publications, followed by medRxiv. Documents indexed in BioRxiv receive on average, the highest social media attention.
7. We expect that the total number of COVID-19 related publications will double in 20 days. Journal articles will double in 24 days, while papers in repositories will grow at a slower rate.
8. We expect non OA papers to grow at a faster rate than OA publications. By mid-May the number of non OA papers will have almost tripled.

## Acknowledgements

This work is supported by the Ministerio español de Economía y Competitividad under project TIN2017-85727-C4-2-P (UGR-DeepBio). Nicolas Robinson-Garcia has received funding from the European Union’s Horizon 2020 research and innovation programme under the Marie Skłodowska-Curie grant agreement No 707404. Daniel Torres-Salinas has received funding from the University of Granada’ Plan Propio de Investigación y Transferencia under the Reincorporación de Jóvenes Doctores grant.

## Footnotes

1 https://pages.semanticscholar.org/coronavirus-research

2 https://www.who.int/emergencies/diseases/novel-coronavirus-2019/global-research-on-novel-coronavirus-2019-ncov

3 Additional information on this dataset is available at https://covid-19.dimensions.ai/.

